# Genetic determinants of chromatin accessibility in T cell activation across humans

**DOI:** 10.1101/090241

**Authors:** Rachel E. Gate, Christine S. Cheng, Aviva P. Aiden, Atsede Siba, Marcin Tabaka, Dmytro Lituiev, Ido Machol, M. Grace Gordon, Meena Subramaniam, Muhammad Shamim, Kendrick L. Hougen, Ivo Wortman, Su-Chen Huang, Neva C. Durand, Ting Feng, Philip L. De Jager, Howard Y. Chang, Erez Lieberman Aiden, Christophe Benoist, Michael A. Beer, Chun J. Ye, Aviv Regev

## Abstract

Over 90% of genetic variants associated with complex human traits map to non-coding regions, but little is understood about how they modulate gene regulation in health and disease. One possible mechanism is that genetic variants affect the activity of one or more *cis*-regulatory elements leading to gene expression variation in specific cell types. To identify such cases, we analyzed Assay for Transposase-Accessible Chromatin sequencing (ATAC-seq) and RNA-seq profiles from activated CD4^+^ T cells of up to 105 healthy donors. We found that regions of accessible chromatin (ATAC-peaks) are co-accessible at kilobase and megabase resolution, in patterns consistent with the 3D organization of chromosomes measured by *in situ* Hi-C in T cells. 15% of genetic variants located within ATAC-peaks affected the accessibility of the corresponding peak through disrupting binding sites for transcription factors important for T cell differentiation and activation. These ATAC quantitative trait nucleotides (ATAC-QTNs) have the largest effects on co-accessible peaks, are associated with gene expression from the same aliquot of cells, are rarely affecting core binding motifs, and are enriched for autoimmune disease variants. Our results provide insights into how natural genetic variants modulate *cis*- regulatory elements, in isolation or in concert, to influence gene expression in primary immune cells that play a key role in many human diseases.

## Introduction

The vast majority of disease-associated loci identified through genome-wide association studies (GWAS)^1-3^ are located in non-coding regions of the genome, often distant from the nearest gene^4^. Quantitative trait loci (QTL) studies that associate genetic variants with molecular traits provide a framework for assessing the gene regulatory potential of disease-associated variants. For example, a statistically significant number of GWAS loci are associated with gene expression (expression QTLs – eQTLs) across diverse cell types and states^5-10^, implicating a role for gene regulation in determining disease risk^11,12^. However, because of linkage disequilibrium and the context specificity of most genetic effects^13,14^, it remains difficult to pinpoint disease-causing variants and to determine the mechanistic basis by which they influence gene expression and increase disease risk.

Genetic analysis of variation in chromatin state^13-17^ is a powerful approach for identifying single nucleotide polymorphisms (SNPs) that directly affect *cis*-regulatory activity^18^. In lymphoblastoid cell lines, thousands of SNPs have been associated with DNase I hypersensitivity (measured by DNase-seq)^19^ and histone tail modifications (measured by ChIP-seq)^20-22^. Similar findings have been reported associating SNPs with variation in DNA methylation and histone tail modifications in primary immune cell types (neutrophils, monocytes and CD4^+^/CD45RA^+^ effector memory T cells)^12^. Furthermore, most of these studies found SNPs associated with variation in chromatin states were also associated with nearby transcript abundance, suggesting the genetic perturbation of *cis*-regulatory activity as a major determinant of gene expression variability^11,12,19,21,23^.

These studies have provided foundational resources for understanding the genetic basis of gene regulation at baseline, but many disease states are associated with cell activation, especially in immune cells^24,25^. In particular, dysregulation of T cell homeostasis and activation are known to play a role in autoimmune diseases^26,27^, cancers^28,29^ and infectious diseases^30^, and hundreds of SNPs have been associated with gene expression during T cell activation and polarization^10,31^. Moreover, both DNase-seq and ChIP-seq are laborious and require large cell numbers, it remains challenging to apply them to primary human cells at the scale required for genetic association. Assay for Transposase-Accessible Chromatin sequencing (ATAC-seq), a simple and efficient two-step protocol for measuring chromatin accessibility with low cell number requirements^32^, enables profiling of chromatin state in disease-relevant primary cells isolated from large human cohorts.

Here, we performed ATAC-seq on activated CD4^+^ T cells from 105 healthy individuals to characterize the extent of natural variability in chromatin state, identify its genetic basis, and assess its influence on gene expression. We further leverage the variability between individuals to identify co-accessible regions of accessible chromatin and to relate those to genetic variation and 3D genome organization in T cells. Our work helps lay the foundation for the critical tasks of dissecting gene regulatory relationships between *cis*-regulatory elements in primary human T cells and characterizing how genetic variation contribute to gene expression variability between individual humans.

## Results

### Changes in T cell chromatin state in response to activation

We used ATAC-seq^32^ to assay CD4^+^ T cells in two different conditions: either unstimulated (Th), or stimulated *in vitro* using tetrameric antibodies against CD3 and CD28 for 48 hours (Th_stim_) (**Fig. 1a**). Aligned reads from six samples (five donors, one pair of replicates) were pooled for each condition, yielding a total of 209 million reads for Th_stim_ and 58 million for Th cells (**Methods**). Of these five donors, two are of East Asian, two are of African American and one is of European decent (**Supplementary Fig. 1**). There was a global increase in chromatin accessibility in response to stimulation, with 52,154 chromatin accessible peaks detected in Th_stim_ (average width of 483 bp +/- 344 bp) and 36,487 in Th cells (average width of 520 bp +/-319 bp) (MACS2, FDR < 0.05). Downsampling each Th_stim_ sample to the same number of reads as the matching Th sample yielded a similar trend of peaks (24,665 Th_stim_ vs. 17,313 Th peaks) suggesting the increased accessibility is not due to differences in sequencing depth. Of the 63,763 peaks identified in at least one condition, 27,446 are similarly accessible between the conditions (shared peaks), 28,017 are more accessible in Th_stim_ cells (Th_stim_-specific peaks) (FDR < 0.05), and only 8,298 are more accessible in Th cells (Th-specific peaks) (FDR < 0.05) (**Fig. 1b** and **Supplementary Table 1**).

**Figure 1.**
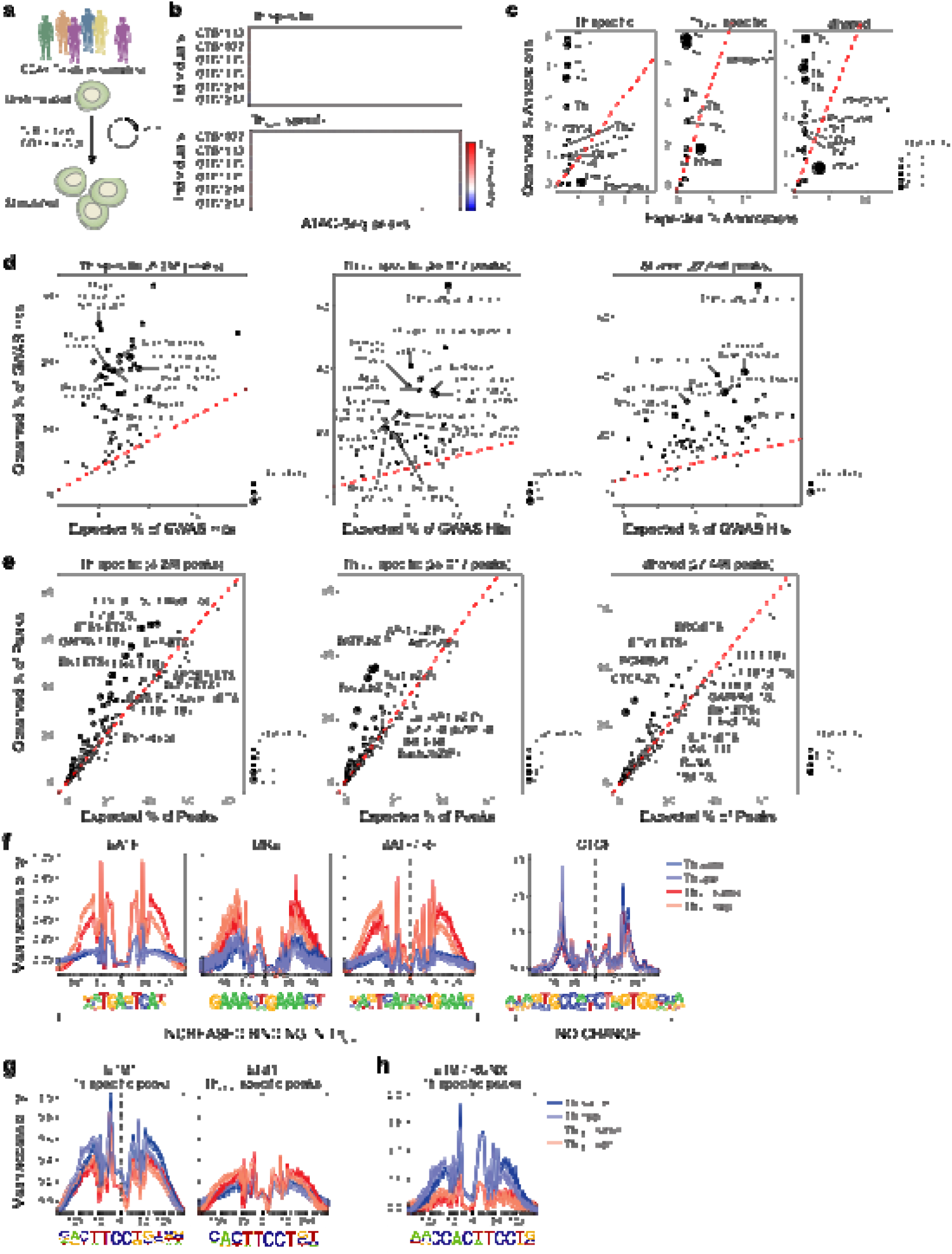
Chromatin dynamics in human T cell activation. **(a)** Experimental overview of ATAC-seq response to activation study. **(b)** Differential chromatin accessibility. Regions of open chromatin (columns) in six samples (rows) before (top, Th specific) and 48hr after (bottom, Th_stim_ specific) activation of primary T cells with anti-CD3/CD28 antibodies. **(c)** Overlap with Th cell enhancers. For each Th annotation, expected (x-axis) versus observed (y-axis) percentage of annotated features overlapping Th-specific (left), Th_stim_-specific (center) and shared peaks (right). **(d)** Overlap with GWAS variants. For each phenotype or disease, expected (x-axis) vs. observed (y-axis) percentage of GWAS loci overlapping Th-specific (left), Th_stim_-specific (center), or shared (right) peaks. **(e)** Transcription factor motif enrichment. Expected (x-axis) vs observed (y-axis) percentage of Th-specific (left), Th_stim_-specific (center), or shared (right) peaks overlapping each TF binding site annotation. **(f-h)** TF footprinting. For each TF motif (as defined in ENCODE^57^), nucleotide resolution average chromatin accessibility (y-axis) in Th (purple) or Th_stim_ (red) cells along the TF binding site (x-axis; log(bp from center of the TF motif)). Aggregated locations are defined as **(f)** Th_stim_-specific peaks overlapping BATF, ISRE, and BATF/IRF motifs (three left panels) and shared peaks overlapping CTCF binding sites (right panel), **(g)** Th-specific (left) and Th_stim_-specific (right) peaks overlapping ETS1 binding sites, and **(h)** Th-specific peaks overlapping ETS1/RUNX combinatorial binding sites.

Regions of accessible chromatin are associated with distinctive genomic features and enriched for SNPs associated with autoimmune diseases. Compared to Th-specific peaks, Th_stim_-specific peaks overlap a higher percentage of enhancers^18^ (defined by H3K27Ac marks) active in αcCD3/αcCD28-activated (Th_0_, 6.9% *vs*. 2.6%) and phorbol myristate acetate (PMA)-stimulated CD4^+^ T cells (Th_stim_, 7.2% *vs*. 3.6%) and a lower percentage of enhancers active in regulatory (T_reg_, 1.4% *vs*. 4.0%), naïve (T_naive_, 1.2% *vs*. 4.9%) and IL17 producing CD4^+^ T cells (Th_17_, 3.2% *vs*. 4.6%) (**Fig. 1c**)^18^. Th_stim_-specific and shared peaks also overlap a higher percentage of SNPs associated with autoimmune diseases, including inflammatory bowel disease (IBD) (32% and 41% *vs*. 20%) and rheumatoid arthritis (21% and 27% *vs*. 13%) (**Fig. 1d**), highlighting the importance of assessing cells under stimulation.

Analyzing regions of accessible chromatin in aggregate provides estimates of the frequencies and single-nucleotide resolution profiles of transcription factor (TF) binding^32^. Th_stim_-specific peaks are enriched for genomic locations bound by TFs important for CD4^+^ T cell activation or differentiation, including members of the AP-1 super family (*e.g.*, 36% contain a BATF binding site) and interferon regulatory factors (*e.g.*, 15% contain a IRF4 binding site)^33-35^ (**Fig. 1e,f**). Th_stim_-specific peaks overlapping regions bound by both BATF and IRF4 (17.4% of peaks)^34^ reveal a different footprint compared to those overlapping regions bound by only one of the TFs (**Fig. 1f**). Conversely, shared peaks are enriched for regions bound by CTCF and BORIS (encoded by CTCFL), two factors known to maintain chromatin state independent of cell type and state^33-35^ (**Fig. 1e**), and their binding footprints are invariant of condition specificity (**Fig. 1f**). ETS1 binding sites overlapping shared and condition-specific peaks have distinct footprints and binding motifs: we observed the canonical ETS1 motif (5‘-CACTTCCTGT-3’) in shared peaks, a 3’ extended motif (5’-CACTTCCTGT**CA**-3’) in Th-specific peaks, and a T/G → T (5’-CACTTCC**T**GT-3’) substitution at the eighth position in Th_stim_-specific peaks, consistent with sequence motifs found at distal ETS1 binding sites (****Fig. 1g****)^36^. Th-specific peaks are more likely to overlap ETS/RUNX binding sites than shared or Th_stim_-specific peaks (OR = 2.7 and 3.9; Fisher’s exact test, P = 2.2×10^-16^ and P = 2.2×10^-16^, respectively) (****Fig. 1h****), which could be due to an enrichment of Th-specific peaks for T_reg_ enhancers known to be bound by the ETS/RUNX complex^37,38^. An additional 6,102 Th_stim_-specific (6.6% of intergenic regions) and 4,118 shared peaks (4.5% of intergenic regions) were located in non-coding regions previously unannotated by H3K27Ac^18,39^, of which 53.5% and 35.6% overlap known binding sites for transcription factors in the AP-1 super family and IRF family, respectively. These results demonstrate that regions of chromatin accessibility captured by ATAC-seq overlap both known enhancers and TF binding sites important for polarization-independent activation of T cells, consistent with our stimulation protocol, and in aggregate reveal high-resolution footprints distinguishing condition specific and combinatorial transcription factor binding.

### Inter-individual variation reveals co-accessibility patterns at multiple genomic scales

Because Th_stim_-peaks, including shared and Th_stim_-specific peaks, better overlap known T cell enhancers, autoimmune disease loci, and binding sites for TFs important for T cell function, we next characterized the inter-individual variability of chromatin accessibility only in stimulated T cells. We optimized the ATAC-seq protocol to profile activated CD4^+^ T cells (**Methods, Supplementary Fig. 2**) from 105 healthy donors in the ImmVar Consortium^10^, all of European descent (**Fig. 2a, Supplementary Fig. 1**). We obtained a median of 37 million (MAD +/-13 million) reads per sample, from highly complex libraries (**Supplementary Fig. 3**), with low mitochondrial DNA (mtDNA) contamination (on average contamination < 3%). We called peaks using a pool of 4.2 billion merged reads from all 105 individuals, resulting in 167,140 peaks (ATAC-peaks) (MACS2, FDR < 0.05). These included 85.1% of the 52,154 Th_stim_ peaks identified in the initial set of six samples from five individuals (**Fig. 1**) with very similar enrichment for GWAS loci (Pearson R = 0.65) and enhancer elements (Pearson R = 0.88) (**Supplementary Fig. 4**).

**Figure 2.**
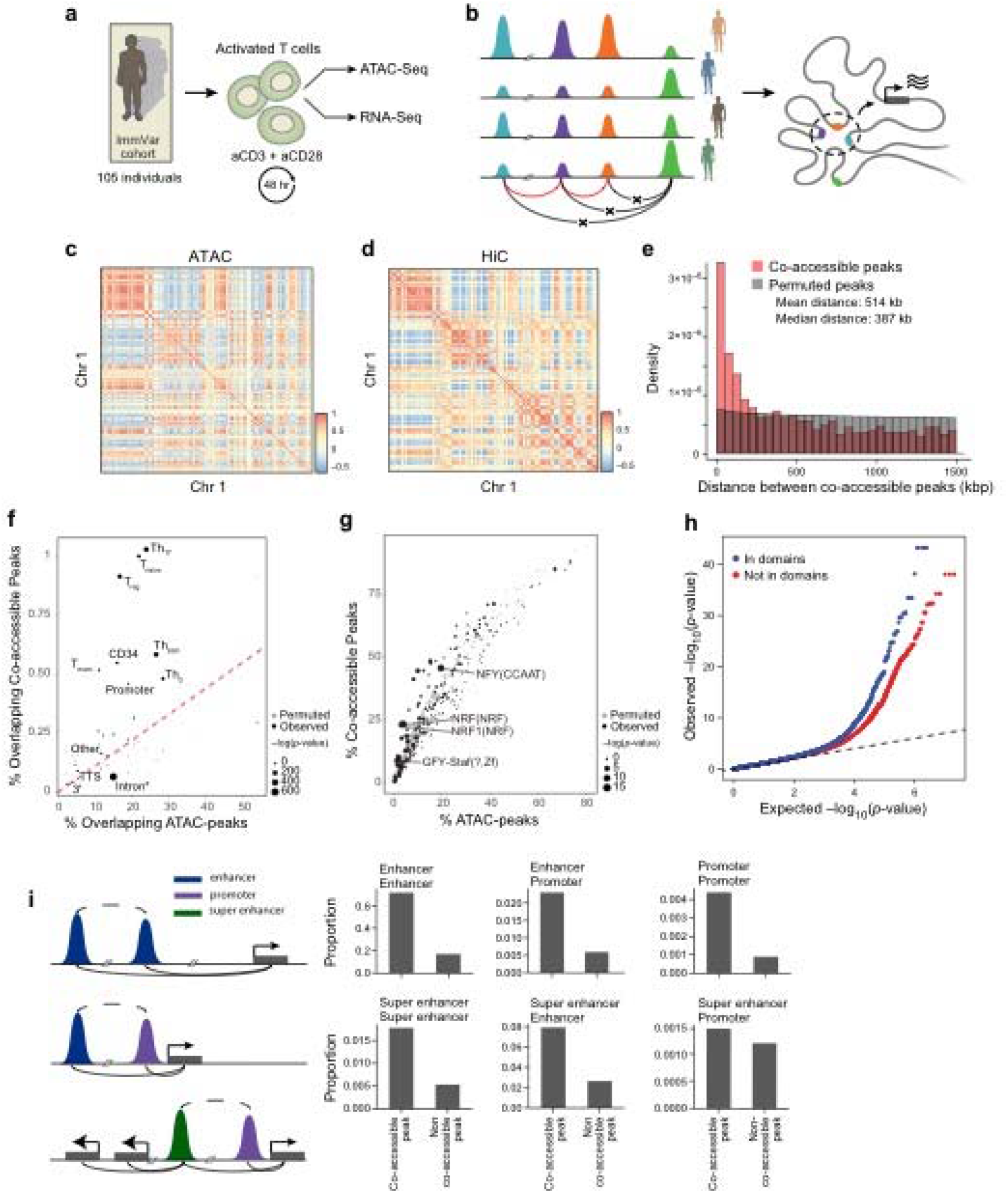
Inter-individual chromatin co-accessibility is constrained by chromosome architecture. **(a)** Overview of T cell activation, ATAC-seq and RNA-seq from 105 samples. **(b)** Cartoon of the multiscale relationship between co-accessible regions across individuals and 3D genome structure. **(c)** Megabase scale inter-individual co-variation of chromatin accessibility. Heat map shows the pairwise Pearson correlation in chromatin accessibility across 105 ATAC-seq profiles in ATAC-peaks binned into 1 Mb windows for Chr 1 (rows, columns). **(d)** Correlation of Hi-C interactions at 1 Mb resolution for Chr 1. **(e)** Length scales of co-accessibility. Histogram of distances between significantly co-accessible peaks (pink) and random permuted peaks (grey). **(f)** Overlap with Th cell enhancers. Percentages of enhancer annotations overlapping all ATAC-peaks (x-axis) versus co-accessible peaks (y-axis). Real peaks (black) and permuted peaks (gray). **(g)** TF motif enrichment. Percentage of ATAC-peaks (x-axis) versus percentage of co-accessible peaks (y-axis) overlapping transcription factor binding sites. Real peaks (black) and permuted peaks (gray). **(h)** Co-accessible peaks overlap with Hi-C domains. Q-Q plot of linear regression p-values for pairs of co-accessible peaks residing in (blue) or out (red) of the same Hi-C domain. **(i)** Pairs of co-accessible peaks overlapping with multiple *cis*-regulatory regions. A cartoon depiction (on left) of co-accessible peaks in enhancer/enhancer (top), enhancer/promoter (middle), and super enhancer/promoter (bottom) regions. Proportion (y-axis) of pairs of co-accessible peaks and non-co-accessible peaks overlapping pairs of annotated *cis*-regulatory elements (on right).

Leveraging the variability in ATAC-peaks across 105 individuals, we found patterns of co-accessibility at multiple genomic scales, recapitulating the 3D conformation of chromatin within chromosomes, as determined by loop-resolution *in situ* Hi-C^40^ of activated CD4^+^ T cells pooled from another five donors (**Fig. 2b, Supplementary Table 2**). At 1 Mb resolution, we observed significant intra-chromosomal co-accessibility, as measured by correlation of total counts of ATAC-seq peaks within each 1 Mb bin (Chr1: **Fig. 2c**, Other chromosomes: **Supplementary Fig. 5**). These pairwise correlations are qualitatively similar to and quantitatively consistent (Pearson R = 0.66) with Hi-C interaction frequencies at the same resolution (**Fig. 2c,d, Supplementary Fig. 5**), likely reflecting variability in the signal (regions of accessible chromatin) to noise (regions of closed chromatin) ratio across samples similar to observations in single cells^32^. At 100 kb resolution, pairwise correlations are also consistent, though to a lesser extent, with Hi-C interaction frequencies (Pearson R = 0.52) (**Supplementary Fig. 6**).

We next characterized peak-resolution chromatin co-accessibility by fitting a linear regression across individuals for every pair of peaks within each 1.5 Mb bin across the genome. After accounting for experimental covariates that affect global patterns of co-accessibility using 15 principal components (**Supplementary Table 3**), we found 2,158 pairs of co-accessible peaks located on average 514 kb apart (permutation, FDR < 0.05, **Fig. 2e, Supplementary Fig. 7, Supplementary Table 3-5**), encompassing 2% (3,204/167,140) of ATAC-peaks. The extent of co-accessible peaks is overall similar to that of all 167,140 ATAC-peaks (**Supplementary Fig. 8a**), but they are individually more likely to overlap T_naïve_, Th_stim_, and Th_17_ enhancers (Fig. 2f) and binding sites for three pioneering factors: NRF, NFY, and STAF (FDR < 0.05, Fig. 2g). Pairs of co-accessible peaks were more correlated when both peaks reside in the same contact domain (estimated from Hi-C interactions, Fig. 2h) and 80% consisted of peaks overlapping pairs of *cis*-regulatory annotations (*e.g.* enhancer/enhancer, enhancer/promoter, super enhancer/promoter; Fig. 2i). Finally, co-accessible peaks were enriched in Th_stim_ super-enhancer regions previously identified^41^ (**Fig. 4c, Supplementary Table 3, Methods**), defined as large clusters of contiguous enhancers, and often bound by master regulators and mediator complexes to drive the transcription of genes involved in cell type specificity^41,42^. These results suggest that co-accessibility between regions of accessible chromatin may be determined by the 3D conformation of the genome and may correspond to coordinated regulation of multiple *cis*-regulatory elements, including known T cell enhancers and regions bound by pioneering factors.

### Genetic variants associated with chromatin accessibility modulate the *cis*-regulatory function of Th cell enhancers

We next defined the genetic basis of chromatin accessibility by associating each ATAC-peak detected in Th_stim_ cells with proximal SNPs across the 105 individuals. To maximize statistical power, we focused only on testing pairs of SNPs (minor allele frequency > 0.05) and ATAC-peaks where the SNP is located *within* the peak, which we term these peaks SNP-containing ATAC-peaks. Of the 64,188 SNP-containing ATAC-peaks tested, 3,318 were significantly associated with a SNP (RASQUAL^43^, *P* < 2.91×10^-3^, permutation FDR < 0.05) (**Fig. 3a, Supplementary Fig. 8b, 9, and Supplementary Table 4-6**). We term each associated SNP a local ATAC quantitative trait nucleotide (*local*-ATAC-QTN) and the corresponding peak a *local*-ATAC-peak. We estimate that 15% of the 64,188 ATAC-peaks are associated with at least one *local*-ATAC-QTN within the peak, using a method to estimate the proportion of null hypotheses while accounting for incomplete power^44^. *Local*-ATAC-peaks exhibited similar read depth as all ATAC-peaks (**Supplementary Fig. 8b**) and the estimates of genetic effects are correlated with those reported using H3K27AC ChIP-seq peaks^12^ (**Supplementary Fig. 10**).

**Figure 3.**
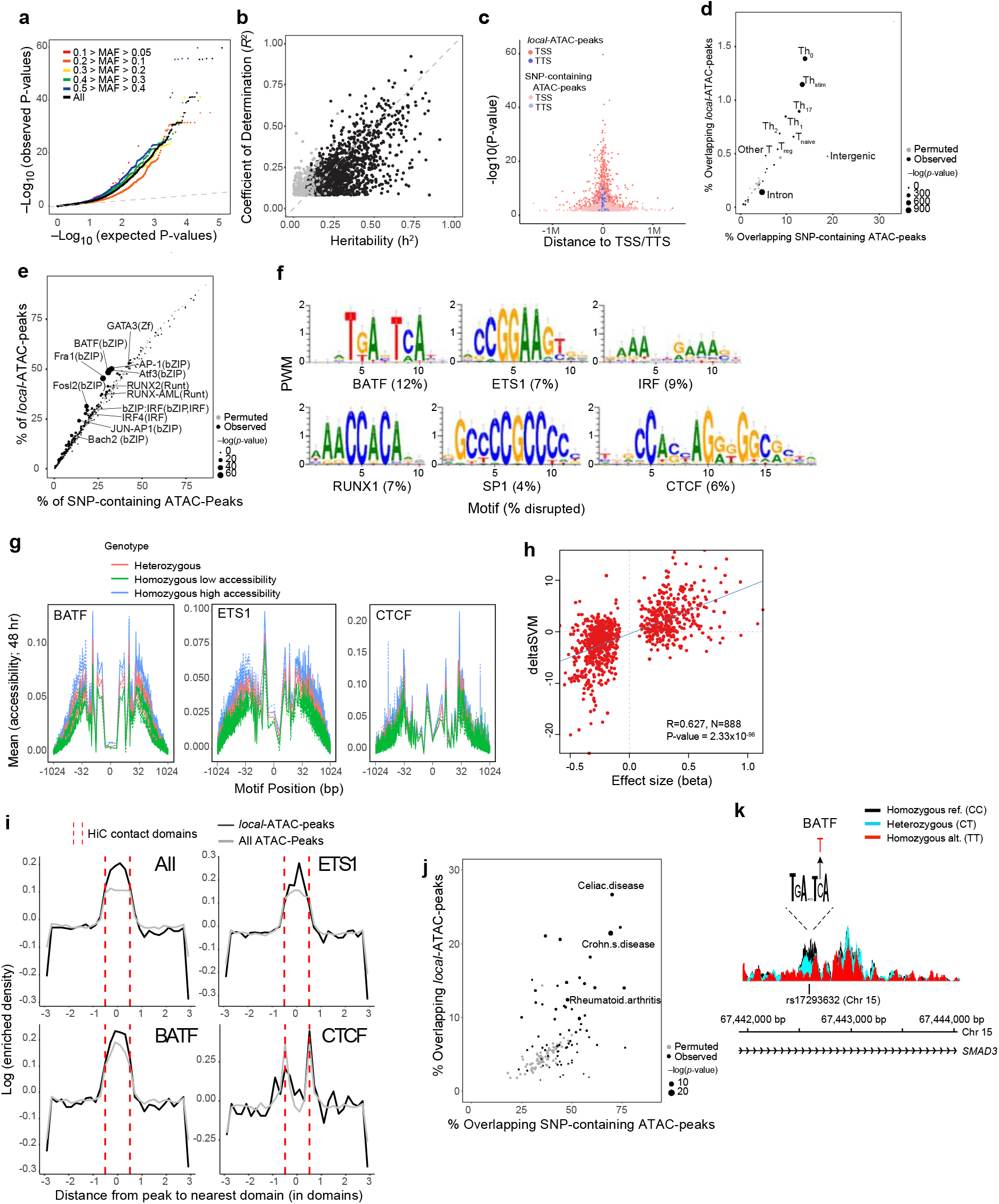
Genetic variants that affect chromatin states in human T cell activation. **(a)** *Local*-ATAC-QTNs. Q-Q plot of linear regression p-values for all *local*-ATAC-QTNs (black) and *local*-ATAC-QTNs filtered for minor allele frequency: 0.1 > MAF > 0.05 (red), 0.2 > MAF > 0.1 (orange), 0.3 > MAF 0.2 (yellow), 0.4 > MAF > 0.3 (green), 0.5 > MAF 0.4 (blue). **(b)** Heritability of chromatin state. For each of 1,428 *local*-ATAC-peaks, coefficient of determination (R^2^) of the best associated *local*-ATAC-QTN (y-axis) versus *cis* heritability (*h*^2^) estimated based on all genotypes +/- 500 kb of each peak (x-axis). Black points: significantly heritable peaks (*q*-value < 0.05). **(c)** Enrichment of *local*-ATAC-peaks in TSS and TTS. For each peak, statistical significance (y-axis) versus the distance of peak to nearest TSS (pink) or TTS (purple). *Local*-ATAC-peaks (dark pink and purple) versus 3,318 randomly sampled SNP-containing ATAC-peaks (light pink and purple). **(d)** Overlap of *local*-ATAC-peaks with Th cell enhancers. Percentage of annotations overlapping SNP-containing ATAC-peaks (x-axis) versus *local*-ATAC-peaks (y-axis). Real peaks (black) and permuted peaks (gray). **(e)** Transcription factor motif enrichment of *local*-ATAC-peaks. Percentage of SNP-containing ATAC-peaks (x-axis) versus percentage of *local*-ATAC-peaks (y-axis) overlapping each TF motif. Real peaks (black) and permuted peaks (gray). **(f-h)** Disruption of TF binding sites by *local*-ATAC-QTNs. **(f)** Unsupervised TF binding site analysis of *local*-ATAC-peaks. Motifs for six TFs associated with most of the large gkmSVM weights, and the percentage of the overall disruption (%, bottom) explained by *local*-ATAC-QTNs. **(g)** Allele specificity of *local*-ATAC-QTNs. For BATF, ETS1 and CTCF motifs (as identified in ENCODE^57^), aggregated plots of mean chromatin accessibility (y-axis) of *local*-ATAC-peaks along the TF binding site (x-axis; log(bp from center of the TF motif)) for samples heterozygous (pink), homozygous for the high (blue) or low (green) *local*-ATAC-QTN alleles. **(h)** Correlation of effect sizes of *local*-ATAC-QTNs **(a)** (x-axis) versus deltaSVM scores (y-axis). **(i)** Relation between contact domains and SNP-containing ATAC-peaks or *local*-ATAC-peaks. For ATAC-peaks or *local*-ATAC-peaks overlapping ETS1, CTCF, or BATF binding sites, enrichment density (y-axis) versus distance (number of domains) of peak to nearest domain (x-axis). Hi-C contact domain boundaries are indicated (dotted red lines). **(j)** Overlap of *local*-ATAC-peaks with GWAS loci. Percentages of GWAS loci overlapping with SNP-containing ATAC-peaks (x-axis) versus *local*-ATAC-peaks (y-axis). Real peaks (black) and permuted peaks (gray). **(k)** Effects of *local*-ATAC-QTN rs17293632 on the accessibility of the corresponding BATF containing *local*-ATAC-peak on chromosome 15. ATAC-seq profiles were aggregated for individuals of different rs17293632 genotypes (black: homozygous major allele, light blue: heterozygous, red: homozygous minor allele).

Several lines of evidence support a model where the accessibility of *local*-ATAC-peaks is determined by *local*-ATAC-QTNs that perturb the function of *cis*-regulatory elements active in stimulated T cells. First, for the 1,428/3,318 *local*-ATAC-peaks that are significantly heritable from fitting a linear mixed model over all SNPs +/- 500kb of each peak (average heritability 44%, GCTA FDR < 0.05), 81% of the heritability is explained by the corresponding *local*-ATAC-QTNs (**Fig. 3b and Supplementary Table 6 and 7; Methods**). This suggests a genetic architecture where a single variant residing in a peak is responsible for the majority of heritable variation. Second, compared to all SNP-containing ATAC-peaks, *local*-ATAC-peaks are preferentially located near transcription start and termination sites (Fig. 3c), are more enriched for T cell enhancers (*P* < 25.76×10^-70^, hypergeometric test; Fig. 3d), and are more enriched for genomic regions bound by TFs involved in T cell development and activation (*e.g.* BATF, AP1 and IRF) (Fig. 3e). Indeed, 72% of *local*-ATAC-peaks contained either a BATF or an ETS1 binding site (1.3-fold enrichment compared to all ATAC-peaks, *P* < 3.39×10^-104^, hypergeometric test) and 29% contained both sites (1.7-fold enrichment, *P* < 1.00×10^-51^, hypergeometric test). Applying a method that predicts the effects of SNPs on TF binding to 903 ATAC-QTNs located within 300 bp of the middle of the corresponding peaks, we found that almost half of ATAC-QTNs (45%) are predicted to strongly disrupt bindings for one of six (BATF, ETS1, IRF, RUNX1, SP1 and CTCF) predicted TF binding sites (TFBSs) (**Fig. 3f**). For *local*-ATAC-peaks that overlap BATF, ETS1 and CTCF binding sites, differential accessibility between genotypes was observed at single nucleotide resolution, with the core motif exhibiting the most striking difference in accessibility, even though only 5% of the corresponding *local*-ATAC-QTNs directly reside in a BATF, ETS1 or CTCF bindings site. This suggests that the perturbation of binding by key TFs – either directly by disrupting their sites, or indirectly by first disrupting binding by other factors in the same regulatory region – may be a major driver for the observed variation in chromatin accessibility across individuals (**Fig. 3g and Supplementary Fig. 11**). An extended 1 kb window exhibited weaker but still significant differences of *local*-ATAC-QTNs on chromatin accessibility (**Fig. 3g and Supplementary Fig. 11**). Consistent with the footprinting analysis, the effect sizes of *local*-ATAC-QTNs are correlated (Pearson R = 0.627, *P* < 2.33×10^-98^) with SNP motif disruption scores obtained by deltaSVM^45^, an unbiased analysis to discover de novo *cis*-regulatory elements in ATAC-peaks (Fig. 3h, Methods). Note that the relation between the accessibility of *local*-ATAC-peaks and 3D chromatin organization is similar to that observed for ATAC-peaks in general (**Fig. 3i**). Both *local*-ATAC-peaks and ATAC-peaks overlapping BATF and ETS motifs are enriched within Hi-C contact domains, whereas those overlapping CTCF motifs are enriched at the contact domain boundaries (Fig. 3i). These results are consistent with previous reports of CTCF enrichment at loop anchors and at contact domain boundaries^40,46-48^.

*Local*-ATAC-peaks are more likely to overlap GWAS SNPs from autoimmune diseases than SNP-containing ATAC-peaks (**Fig. 3j**), providing a functional context for interpreting disease associations. Even though *local*-ATAC-peaks consist of only ~5% of the SNP-containing ATAC-peaks tested, they encompass a large percentage of the loci associated with autoimmune diseases including Celiac‘s disease (28%), Crohn’s disease (22%), and Rheumatoid Arthritis (12%), a 8-fold (hypergeometric *P* < 4.34×10^-7^), 6-fold (hypergeometric *P* < 8.58×10^-17^), and 5-fold (hypergeometric *P* < 6.18×10^-8^) enrichment compared to all SNP-containing ATAC-peaks, respectively. In general, ATAC-peaks encompass 70% of Celiac’s and Crohn’s disease and 47% of Rheumatoid Arthritis GWAS loci. For example, rs17293632 has been associated with Crohn’s disease and IBD^49^ and is located in the first intron of *SMAD3*, a transcription factor involved in the TGF-β signaling pathway that regulates T cell activation and metabolism^50^. This SNP disrupts a consensus BATF binding site at a conserved position (deltaSVM=-12.72), and results in decreased chromatin accessibility in individuals that possess the alternate allele (**Fig. 3k**).

Together, these results suggest that when genetic control of chromatin accessibility is determined by *local*-ATAC-QTNs residing within peaks, this often involves the disruption of binding of key TFs, even though the variants almost always reside outside of the core motif sequence.

Moreover, they show that *local*-ATAC-QTNs overlap a high percentage of autoimmune disease GWAS loci, as in the case of rs17293632.

## Genetic determinants of chromatin co-accessibility

Given the identified genetic determinants of chromatin accessibility and the observed co-accessibility of thousands of peaks, we next tested the hypothesis that *local*-ATAC-QTNs could modulate the accessibility of co-accessible peaks *distally* (within +/- 500kb). We first assessed the heritability of peaks and co-accessible peaks using all SNPs +/- 500kb of each peak. As expected, *local*-ATAC-peaks (2,444/3,318 that converged) were on average more heritable (mean h^2^ = 0.22) than other SNP-overlapping (mean h^2^ = 0.04) and SNP-excluding peaks (mean h^2^ = 0.04) (**Fig. 4a**). Co-accessible peaks, overlapping SNPs or not, were more heritable with a mean heritability of 0.44 and 0.10, respectively. Excluding the 3,318 *local*-ATAC-peaks, we identified 382 that were associated *distally* with a *local*-ATAC-QTN (RASQUAL, *P* < 1.27×10^-4^, permutation FDR < 0.05) located +/- 500 kb from the peak. We term each associated variant a *distal*-ATAC-QTN and each associated peak a *distal*-ATAC-peak. Consistent with the heritability analysis, *distal*-ATAC-QTNs imparted the strongest effects on co-accessible peaks, both residing in or outside of the same Hi-C domain (**Fig. 4b** and **Supplementary Table 2 and 3**).

**Figure 4.**
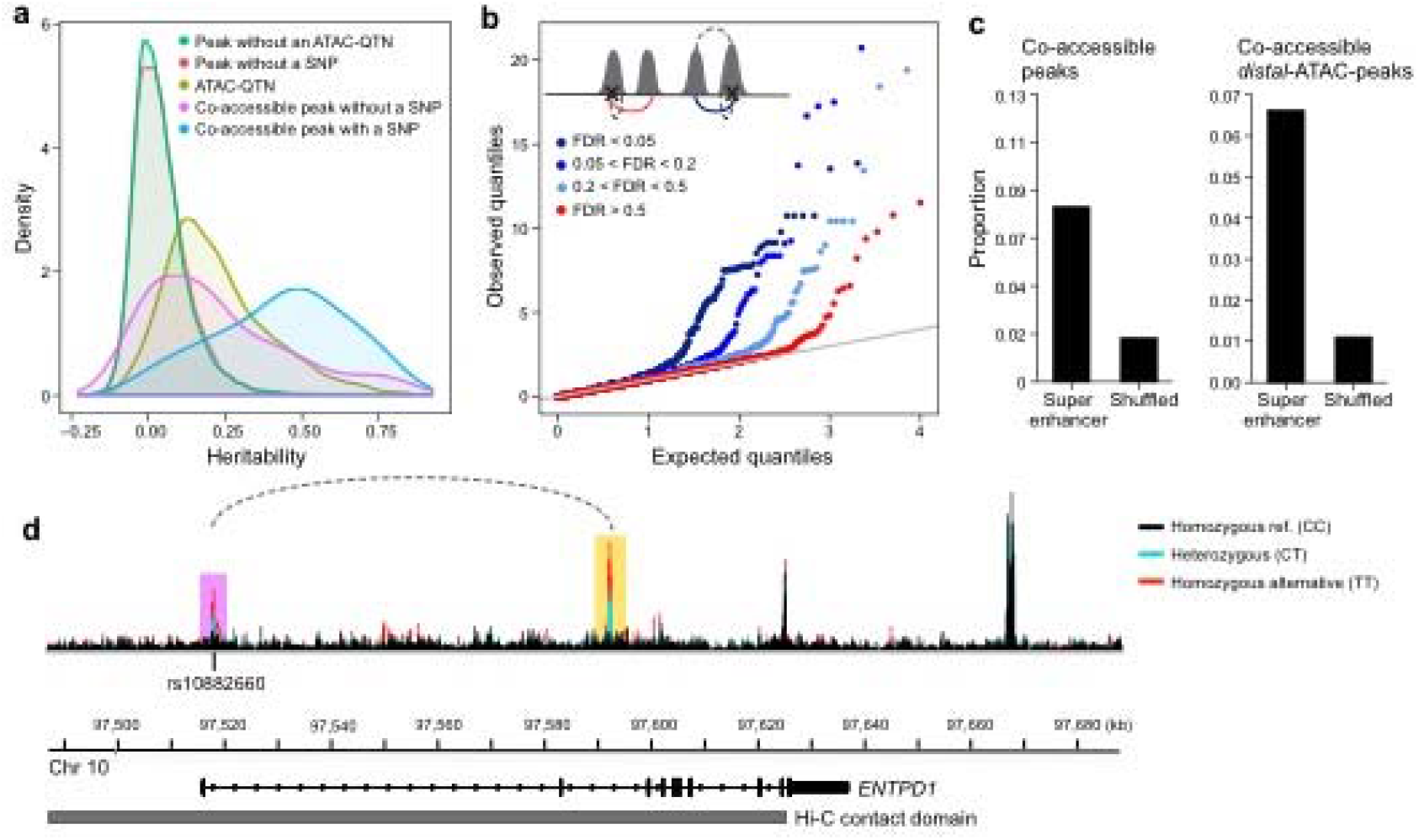
Genetic determinants of co-accessible peaks. **(a)** Distribution of local heritabilities of ATAC-peaks without a *local*-ATAC-QTN (green), ATAC-peaks without a SNP (red), *local*-ATAC-peaks (olive), co-accessible peaks without a SNP (purple), and co-accessible peaks with a SNP (blue). **(b)** Q-Q plots of the linear regression p-values of *distal*-ATAC-peaks that are single peaks (red: co-accessibility FDR > 0.5), or co-accessible peaks at various significance cutoffs (light blue: 0.2 < FDR < 0.5, medium blue: 0.05 < FDR < 0.2, dark blue: FDR < 0.05). The cartoon (upper left corner) depicts the *distal*-ATAC-QTN associations that are single peaks (red line) and co-accessible peaks (blue line). **(c)** Proportion of co-accessible (left), co-accessible *local*-ATAC-peaks (middle) and co-accessible *local*-ATAC-peaks residing in domains (right) overlapping super-enhancers (y-axis) to randomly shuffled super enhancers (x-axis). **(d)** An example of a genetic variant (rs10882660) residing in the first intron in *ENTPD1*, associated locally (in purple) and *distally* (in yellow) to ATAC-peaks. The local and *distal*-ATAC-peaks are co-accessible (dotted line) and reside in a Hi-C contact domain (grey). ATAC-seq profiles were aggregated for individuals of different rs10882660 genotypes (black: homozygous major allele, light blue: heterozygous, red: homozygous minor allele).

Co-accessible peaks and co-accessible *distal*-ATAC-peaks are both more likely to overlap Th_stim_ super enhancers than randomly shuffled super enhancers^41^. The effect is stronger in co-accessible *distal*-ATAC-peaks (6-fold versus 4-fold), suggesting the genetic control of co-accessible peaks likely reflect a shared genetic effect on clusters of enhancers (**Fig. 4c**). In an example, rs10882660 is a local and *distal*-ATAC-QTN for a pair of co-accessible peaks residing in the 1^st^ and 2^nd^ introns of ectonucleoside triphosphate diphosphohydrolase I (*ENTPD1*) that reside in a Hi-C contact domain, likely influencing the coordinated control of these chromatin accessible regions (**Fig. 4d**). *ENTPD1* is one of the dominant drivers of hydrolysis of adenosine triphosphate (ATP) and adenosine diphosphate (ADP) in T_regs_ cells whose expression can lead to tumor growth in mouse models^51-54^. These results and example suggest a model where genetic variation can impart coordinated control on pairs of *cis*-regulatory regions.

### Linking chromatin accessibility to gene expression

We next assessed how *local*-ATAC-QTNs could also influence gene expression. We measured RNA-seq profiles of stimulated CD4^+^ T cells from 95 donors (92 from an aliquot of the same cells with matching ATAC-seq data). We identified 424 genes significantly associated with at least one of 6,903 *local*-ATAC-QTNs located +/- 500 kb from the center of each gene (RASQUAL, *P* < 1.65×10^-3^, permutation FDR < 0.05, **Fig. 5a, Supplementary Tables 4 and 8**). We term the 383 associated SNPs expression quantitative nucleotides (eQTNs) and the corresponding 424 genes eGenes. We estimate that 30% of *local*-ATAC-QTNs are also eQTNs (with a procedure to estimate the proportion of null hypotheses; Methods), consistent with previous reports in LCLs^19,21^. Considering all genetic variants within a 1 Mb region of each eGene, we found 191/424 genes to be highly heritable (GCTA FDR < 0.05), with the eQTN explaining on average 68% of the heritability (**Fig. 5b** and **Supplementary Table 9**). The lower estimates of explained heritability than *local*-ATAC-peaks suggests that the genetic control of gene expression may involve more than one SNP and *cis*-regulatory element in many cases.

**Figure 5.**
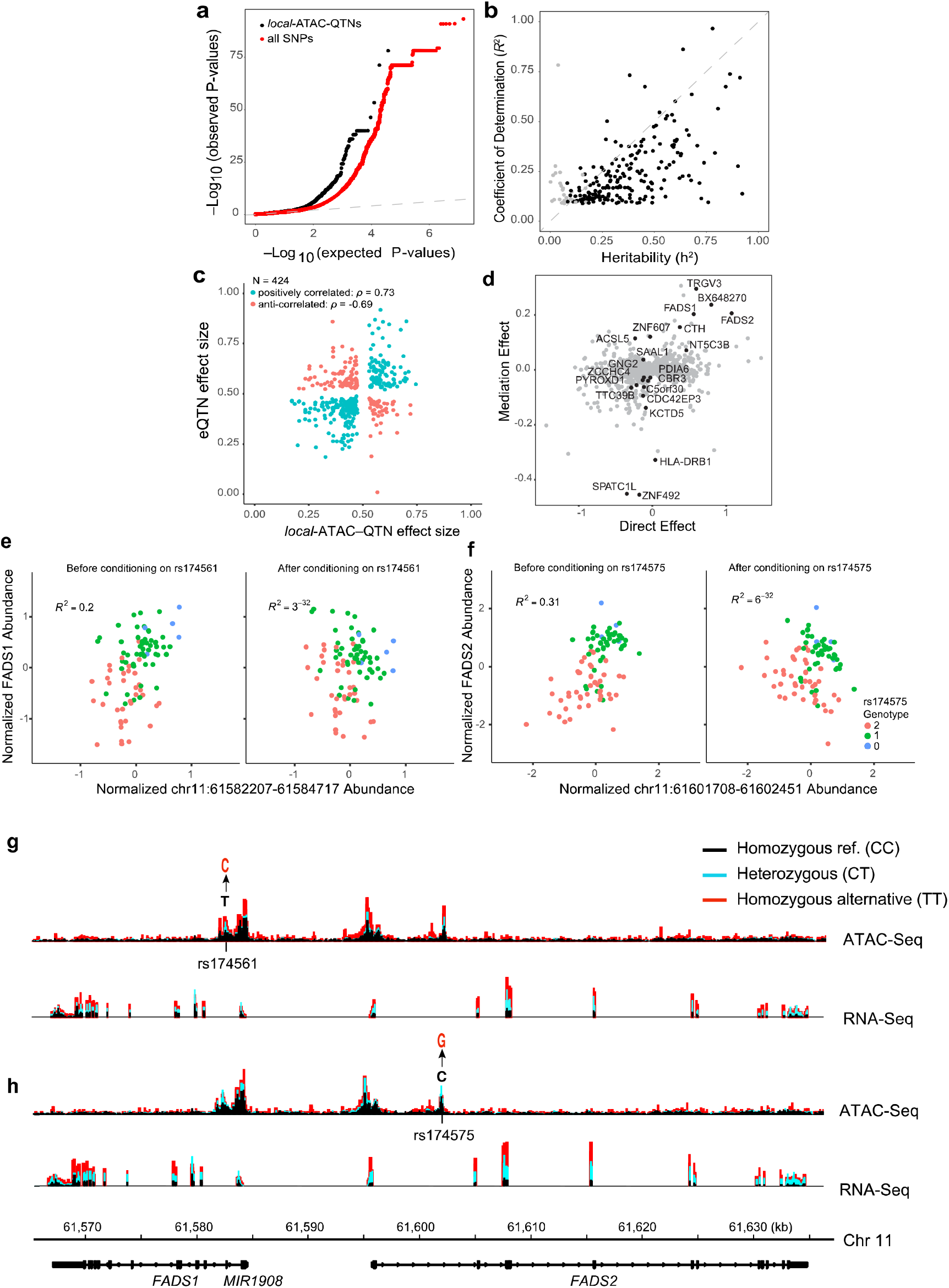
Association of chromatin accessibility and gene expression. **(a)** eQTNs. Q-Q plot of associations between *local*-ATAC-QTNs and expression of genes +/- 500kb. **(b)** Heritability of gene expression. For each of 191 eGenes, coefficient of determination (R^2^) of the best associated eQTN (y-axis) versus *cis* heritability (h^2^) estimated based on all genotypes +/- 500 kb of each gene (x-axis). Black points: significantly heritable peaks (*q*-value < 0.05). **(c)** Correlation of effect sizes between local -ATAC-QTNs (x-axis) and eQTNs (y-axis). **(d)** Mediation of eGenes. Average causal mediation effect estimates (y-axis) and average direct effect estimates (x-axis) for *local*-ATAC-peaks (mediator) and eGenes (outcome variable) sharing a SNP (instrument variable). FDR < 0.1 *local*-ATAC-peaks are colored in black. **(e,f)** Examples of gene expression conditioned on chromatin accessibility. **(e)** *FADS1* expression (y-axis) *vs*. chromatin accessibility at chr11:61582207-61584717 (x-axis) before (left) and after (right) conditioning, colored by rs174561 genotypes. **(f)** *FADS2* expression (y-axis) *vs*. chromatin accessibility at chr11:61601708-61602451 (x-axis) before (left) and after (right) conditioning on rs174575. **(g,h)** ATAC-seq (top) and RNA-seq (bottom) profiles were aggregated for individuals of different **(g)** rs174561 and **(h)** rs174575 genotypes (black: homozygous major allele, light blue: heterozygous, red: homozygous minor allele).

Among the 383 SNPs that are simultaneously associated with chromatin accessibility (as *local*-ATAC-QTNs) and gene expression (as eQTNs), the majority (286/383) have effect sizes in the same direction (Spearman ρ = 0.73) indicative of activating effects, while 138 have effect sizes in the opposite direction indicative of repressive effects (Spearman ρ = -0.69) (**Fig. 5c**).

We next examined the sharing of genetic effects between *local*-ATAC-peaks and eGenes using a bivariate linear mixed model^55^ and mediation analysis^56^. Because of limited sample size, measuring the genetic correlation for individual pairs of *local*-ATAC-peaks and eGenes is likely under powered. However, the distribution of genetic correlations for 161 pairs of *local*-ATAC-peaks and eGenes that converged (inverse variance weighted average of 0.66) was statistically different from both randomly sampled (inverse variance weighted average of 0.23, Kolmogorov-Smirnov *P* = 4.32×10^-10^) and permuted ATAC-peaks (inverse variance weighted average of 0.07, Kolmogorov-Smirnov *P* = 1.68×10^-10^) (**Supplementary Fig. 12** and **Supplementary Table 10**). This is corroborated by mediation analysis where the genetic effects on 21/424 eGenes were significantly mediated by the corresponding *local*-ATAC-peaks (FDR < 0.1, **Fig. 5d**) and the high correlation of the mediation effects and the inverse variance weighted genetic correlation (Pearson R = 0.52, *P* < 1.2×10^-12^, **Supplementary Fig. 13**). For example, before conditioning on rs174575, *FADS2* expression and chromatin accessibility at chr11:61,601,708-61,602,451 are correlated (R^2^ = 0.31, *P* < 8.25×10^-9^) and after conditioning the correlation is no longer significant (R^2^ = 6×10^-32^, *P* < 1). Similarly, after conditioning on rs174561, *FADS1* is no longer correlated to chr11:61,601,708-61,602,451 (before conditioning: R^2^ = 0.2, *P* < 8.74×10^-6^; after conditioning: R^2^ = 3×10^-32^, *P* < 1) (**Fig. 5e,f**). Notably, rs174561 is a *local*-ATAC-QTN associated with a pair of co-accessible peaks and is a variant previously associated with Crohn’s disease^57^ (**Fig. 5f**). The associated co-accessible peaks span the promoters of *FADS1* and *FADS2*, which are two fatty acid desaturases (FADS) that regulate inflammation, promote cancer development, and *FADS2* knockout mice develop dermal and intestinal ulcerations (**Fig. 5g,h**)^58-61^. These results suggest that variability in chromatin accessibility may underlie and mediate variability in gene expression and, at the *FADS1* and *FADS2* loci, increase risk for Crohn’s disease.

## Discussion

Variability in gene expression is a hallmark of all naturally segregating populations and has been extensively characterized in cell lines and primary cells. Variability in chromatin state, on the other hand, has been challenging to measure in primary cells due to the lack of scalable assays that could operate on limited cell numbers. Toward this end, we analyzed ATAC-Seq profiles from five individuals at both baseline and after stimulation. We found substantial remodeling of chromatin organization, with a significantly higher number of accessible regions, overlapping a substantially higher proportion of SNPs associated with autoimmune diseases. This suggests the importance of exploring this relationship in the context of specific, physiologically-relevant stimulations. Notably, regions open under stimulation overlap different enhancer subsets suggesting that analysis of different Th cell polarizations (T_regs_, Th_17_ etc.) could illuminate additional relevant events.

Based on this rationale, we then dissected the relationship between genetic variation and variation in chromatin accessibility in stimulated Th cells from 105 individuals. Variation across individuals highlights four inter-related phenomena. **First**, accessible regions co-vary across the genome of an individual (co-accessibility) at multiple length scales, reflective of the 3D structure of the genome. At individual peak resolution, ~2% of the peaks are co-accessible, especially if they are within the same 3D contact domain, and these are more likely to overlap Th enhancers and binding sites for T cell pioneering factors, and consist of “pairs” of regulatory regions, including super-enhancers. These results suggest that co-accessibility between pairs of peaks may be determined by the 3D conformation of the genome and may correspond to coordinated regulation of multiple *cis*-regulatory elements, especially by pioneering factors. The limited co-accessibility is consistent with results from analyzing DNase-I hypersensitivity regions but lower than those observed from analyzing histone ChIP-seq marks. As previously suggested, this could be due to either differences in the types of genomic interactions captured by each assay or depth of sequencing between studies. **Second**, combining genetic variation with variation in individual peak accessibility, we identified *local*-ATAC-QTNs: SNPs within an ATAC-peak that explain its variation across individuals. Even though only a minority (5%) of *local*-ATAC-QTNs directly reside within the core binding sites of pioneer TFs, nearly half (45%) are predicted to dramatically disrupt binding at TF sites, based on their allele-specific impact on accessibility. Moreover, even though *local*-ATAC-peaks are only 5% of SNP-containing ATAC-peaks, they overlap ~10^-30^% of the loci for several common autoimmune diseases. The overwhelming enrichment for autoimmune disease loci among *local*-ATAC-peaks could be the result of both the increased number of features tracking cell state and the propensity for disease-causing variants to perturb *cis*-regulatory elements containing key TFs active in specific cell types or states, such as effector or regulatory T cells. **Third**, we found that *local*-ATAC-QTNs can further act *distally* on additional peaks in a 1.5 Mb window, with the strongest effects on those local and *distal*-ATAC-peaks that are co-accessible, which is likely to substantially increase their mechanistic and functional impact. **Fourth**, considering *local*-ATAC-QTNs in the context of variation in gene expression (by RNA-Seq of 92 shared individuals), we estimated that 30% of *local*-ATAC-QTNs are also eQTNs, with bivariate and mediation analyses supporting mechanistic directionality in variability in chromatin accessibility and gene expression.

In a manner consistent with known modes of transcriptional regulation, our approach for a staged analysis, testing the effects of *local*-ATAC-QTNs on *distal*-ATAC-peaks and gene expression, allowed us to overcome power limitations from the sample size and the technical and biological variability in the assays to detect hundreds of genes associated with *local*-ATAC-QTNs. Despite this, there was limited power for bivariate analysis to quantify the shared genetic effects and establish causality for the observed association to both chromatin state and gene expression. These limitations will likely be overcome in future studies with larger sample sizes and higher sequencing depth.

Our findings, derived from large scale genetic association of quantitative chromatin and gene expression traits in primary human cells implicated in many diseases, provide a molecular framework for how disease-causing variants could alter local chromatin structure to modulate gene expression. With the recent advancement of single cell epigenomic^62^ and transcriptomic^63,64^ profiling, it should be possible to more directly detect context-specific genetic effects in a heterogeneous cell population. Future studies that use other disease-relevant primary cells and tissues will help pinpoint causal disease variants and understand the regulatory mechanism underlying common disease.

## Materials and Methods

### Study subjects and genotyping

Healthy subjects between the ages of 18 to 56 (avg. 29.9) enrolled in the PhenoGenetic study^8^ were recruited from the Greater Boston Area and gave written informed consent for the studies. Individuals were excluded if they had a history of inflammatory disease, autoimmune disease, chronic metabolic disorders or chronic infectious disorders. Genotyping using the Illumina Infinium Human OmniExpress Exome BeadChips (704,808 SNPs are common variants [MAF > 0.01] and 246,229 are part of the exomes; Illumina Inc., San Diego, CA) has been previously described^18^. The genotype success rate was at least 97%. We applied rigorous subject and SNP quality control (QC) that includes: (1) gender misidentification; (2) subject relatedness; (3) Hardy-Weinberg Equilibrium testing; (4) use concordance to infer SNP quality; (5) genotype call rate; (6) heterozygosity outlier; and (7) subject mismatches. We excluded 1,987 SNPs with a call rate < 95%, 459 SNPs with Hardy-Weinberg equilibrium p-value < 10^-6^, and 63,781 SNPs with MAF < 1% from the 704,808 common SNPs (a total of 66,461 SNPs excluded). Principal component analysis of genotypes from all individuals used in the study are shown in **Supplementary Figure S6**.

We used the IMPUTE2 software (version: 2.3.2) to impute the post-QC genotyped markers from the entire Immvar cohort (N = 688) using reference haplotype panels from the 1000 Genomes Project (The 1000 Genomes Project Consortium Phase III) that contain a total of 37.9 Million SNPs in 2,504 individuals with ancestries from West Africa, East Asia, and Europe. After genotype imputation, we extracted the genotypes for 105 individuals assayed for chromatin accessibility and gene expression. Additional filtering for SNPs with MAF < 0.05 in our cohort resulted in 4,558,693 and 4,421,936 common variants tested for chromatin accessibility and gene expression assays, respectively.

### Preparation and activation of primary human CD4^+^ T cells

CD4^+^ T cells were isolated and stimulated as previously described^10^. Briefly, CD4^+^ T cells were isolated from whole blood by negative selection using RosetteSep human CD4^+^ T cell enrichment cocktail (STEMCELL Technologies Inc., Vancouver, BC) and RosetteSep density medium gradient centrifugation. Isolated CD4^+^ T cells were placed in freezing container at -80°C for overnight, and then moved into a liquid nitrogen tank for long-term storage. On the day of activation, CD4^+^ T cells were thawed in a 37°C water bath, counted and resuspended in RPMI-1640 supplemented with 10% FCS, and plated at 50,000 cells per well in a 96 well round-bottom plate. Cells were either left untreated or stimulated with beads conjugated with anti-CD3 and anti-CD^28^ antibodies (Dynabeads, Invitrogen #11131D, Life Technologies) at a cell:bead ratio of 1:1 for 48 hours, a time point we previously found to maximize the gene expression response in CD4^+^ T cells. At each time point, cells were further purified by a second step positive selection with CD4^+^ Dynabeads (Invitrogen #11145D, Life Technologies).

### ATAC-seq profiling

ATAC-seq profiles were collected for 139 individuals (**Supplementary Table 4**). We performed ATAC-seq as previously described^32^, with a modification in the lysis buffer to reduce mitochondrial DNA contamination, while maintaining high complexity of nuclear reads. 200,000 purified CD4^+^ T cells were lysed with cold lysis buffer (10 mM Tris-HCl, pH 7.4, 10 mM NaCl, 3 mM MgCl2 and 0.03% tween^20^). Immediately after lysis, nuclei were spun at 500g for 8 minutes at 4°C. After pelleting the nuclei, we carefully removed the supernatant and resuspended the nuclei with Tn5 transposase reaction mix (25 ul 2X TD buffer, 2.5 ul Tn5 transposase, and 22.5 ul nuclease-free water) (Illumina Inc). The transposition reaction was performed at 37°C for 30 minutes. Immediately after the transposition reaction, DNA was purified using a Qiagen MinElute kit. Libraries were sequenced on an Illumina HiSeq 2500 sequencer to an average read depth of 42 million (+/- 38 million) per sample (**Supplementary Fig. S2**), with low mtDNA contamination (0.30 – 5.39%, 1.96% on average), low rates of multiply mapped reads (6.7 – 56%, 19% on average) and a relatively high percentage of usable nuclear reads (60 – 92%, 79% on average).

### RNA-seq profiling

RNA-seq profiles were collected for 95 individuals, of which 93 have matching ATAC-seq profiles (**Supplementary Table 4**). RNA was isolated using Qiagen RNeasy Plus Mini Kit and RNA integrity was quantified by Agilent RNA 6000 Nano Kit using the Agilent Bioanalyzer. Purified RNA were converted to RNA-seq libraries using a previously published protocol^65^, where reverse transcription was carried out based on the SMART template switching method and the resulting cDNA was further tagmented and PCR amplified using Nextera XT DNA Sample kit (Illumina) to add the Illumina sequencing adaptors. Samples were sequenced on Illumina HiSeq 2500 to an average depth of 16.9 million reads per sample (+/- 8.7 million).

### *In situ* Hi-C

CD4^+^ T cells were isolated from commercially available fresh blood of healthy individuals (Research Blood Components). CD4^+^ T cells were stimulated for 48 hours with beads conjugated with anti-CD3 and anti-CD^28^ antibodies. *In situ* Hi-C was performed as previously described^40^. Cells were crosslinked with 1% formaldehyde for 10 min at room temperature. After nuclei permeabilization, DNA was digested with MboI and digested fragments were labeled using biotinylated d-ATP and ligated. After reverse crosslinking, ligated DNA was purified and sheared to ~400 bp. Biotin labeled DNA fragments were then pulled down with streptavidin beads and prepped for Illumina sequencing^40^. The final libraries were sequenced using Illumina HiSeq and NextSeq to produce ~3.5 billion 100bp paired-end reads.

### Alignment of ATAC-seq reads

25bp ATAC-seq reads were aligned to the human genome assembly (hg19) with the Burrows Wheeler Aligner-MEM (version: 0.7.12)66. For each sample, mitochondrial reads were filtered out using BEDtools (function intersectBed) and multiply-mapped reads were filtered using Samtools “view” with option “-F 4”67. After filtering, we had a median of 37 million (MAD +/-13 million) reads per sample.

### ATAC-seq peak identification

Filtered ATAC-seq reads from six matched samples (5 individuals and 1 biological replicate) for Th and Th_stim_ cells were merged (separately for Th and Th_stim_ cells) using the Samtools function “merge”. Peaks were called on the respective Th and Th_stim_ merged bam files using MACS2 – callpeak (with parameters -nomodel, -extsize 200, and -shift 100), resulting in 36,486 Th peaks with an average width of 520 bp (+/- 319 bp) and 52,154 Th_stim_ peaks with an average width of 483 bp (+/- 344 bp) (Benjamini-Hochberg FDR < 0.05). The Th and Th_stim_ peaks were further merged (using the BEDtools “merge” function), to a total of 63,763 jointly called peaks. A matrix of the coverage for each of the 63,763 peaks in each of the 12 samples was used as input for detecting differentially accessible peaks. Differentially accessible peaks between Th and Th_stim_ conditions were identified using the DESeq2 R package (version 3.2)68, with 8,298 Th-specific peaks (FDR < 0.05, more accessibility in Th cells), 28,017 Th_stim_-specific peaks (FDR < 0.05, more accessibility in Th_stim_), and 27,446 shared peaks (FDR > 0.05).

For the co-accessibility and genetic analyses, 4.2 billion filtered ATAC-seq reads from 105 Th_stim_ samples were merged to call 167,140 peaks (FDR < 0.05) as previously stated, at an average peak size of 642 bp (+/- 512 bp). Coverage for each peak over all 105 samples was computed to generate a 167,140 peaks x 105 individuals matrix that was used for co-accessibility peak analysis and genetic analyses.

### Percentage of peaks overlapping transcription factor binding motifs

We used the Homer suite, which uses ChIP-seq data from the ENCODE^69^ and Epigenomics Roadmap^14^ projects, to determine transcription factor enrichment within our ATAC-peaks. Percentages of MACS2 called peaks overlapping TF binding motifs were computed using the default setting in the function findMotifsGenome.pl (with parameters hg19, –size given), using the Homer default background. For co-accessible peaks and *local*-ATAC-peaks, peaks were randomly permuted, while retaining the width of each peak to assess the expected TF motif enrichment.

### Transcription factor footprinting

Using the Homer suite tool annotatePeaks, and options –m and –mbed, we found all instances of BATF, ISRE, BATF/IRF, ETS1, and CTCF motifs in shared, Th-specific and Th_stim_-specific peaks. Next, we determined the per-base coverage +/- 1 kb around the center of the motif, only using cut-site reads and splitting the reads into those that map to the same or opposite strand as the motif. For each TF footprint, we generated a matrix with the number of rows equal to the number of instances of the motif by 4,000 columns quantifying coverage: upstream 1 kb from the same strand as the motif, downstream 1 kb from the same strand as the motif, upstream 1 kb from the opposite strand as the motif, downstream 1 kb from the opposite strand as the motif. Final TF footprints were derived from median normalized reads^69^.

### Outlier analysis and sample mix-up analysis

We kept samples that were highly correlated for downstream analyses (Pearson R > 0.68, **SOM**), where we developed an optimized ATAC-seq protocol (**SOM**) that achieved high technical and biological reproducibility (**Supplementary Fig. S1**), highly complex libraries (on average 84% usable nuclear reads, as opposed to 40% prior to optimization) (**Supplementary Fig. S2**), and low mitochrondrial DNA (mtDNA) contamination (on average contamination < 3%, as opposed to 53% prior to optimization). Quality of all ATAC-seq samples were assessed again, only keeping samples that contain a minimum of 8 million QC-passed reads (median of 37 million, MAD +/-13 million) and high inter-sample correlation (Pearson R > 0.68, **SOM**). ATAC-Seq profiles from the 105 individuals were further filtered to identify sample mix-ups. We used the software VerifyBamID^70^ to match each ATAC-seq and RNA-seq sample with the genotyping profile with the highest fIBD score. Samples with designated labels not matching the VerifyBamID predicted genotyping labels were flagged as sample mix-ups. We switched the designated label to the predicted label for cases where the fIBD > 90%. 15 out of the 139 total ATAC-seq samples were re-labeled and 4 out of the 110 total RNA-seq samples were re-labeled. For the ATAC-seq samples: 18 do not have genotypes, 3 are outliers, 1 did not match anyone. For the 110 RNA-seq samples: 8 samples do not have genotypes, 5 are outliers, 1 did not match anyone. 111 ATAC-seq samples and 96 RNA-seq samples were used in the final analysis after filtering for outliers (average mean correlation to others samples < 0.7). In the response to activation study, there were 5 people total, 1 person was repeated for a total of 6 samples, none were genotyped.

### Genetic association analysis of ATAC-QTNs

Genetic association analysis was performed on 105 samples of European descent (**Supplementary Fig. S6**) by running RASQUAL^43^ on the 167,140 peaks identified in Th_stim_ cells and 4,558,693 imputed genetic variants, testing variants within a 1 Mb window of each ATAC-peak, and filtering for a minor allele frequency of greater than 5%. The input to RASQUAL is the number of raw reads in each peak quantified using BEDtools “coverage” using uniquely mapped nuclear reads from each of the 105 individuals. Normalization of reads is performed internal to RASQUAL as previously described [ref RASQUAL]. Duplicated fragments were kept for quantification. Sex and ten principal components were included as covariates to minimize the effects of confounding factors. Using the RASQUAL “-r” option, 10 random permutations for each ATAC-peak were generated. For local -ATAC-peak analysis, only observed and permuted association statistics for peak-SNP pairs where the SNP resides within the peak are compared. For *distal*-ATAC-peak analysis, observed and permuted association statistics for peak-SNP pairs where the SNP does not reside within the peak are compared. In each case, empirical P-values were computed using the R qvalue^44^ package to detect a total of 3,318 local -ATAC-peaks (FDR < 0.05) and 2,376 *distal*-ATAC-peaks (FDR < 0.05).

### Hi-C data analysis

The sequenced reads were analyzed using the Juicer pipeline^71^. We sequenced 402,533,981 Hi-C read pairs in unstimulated T cells yielding 704,323,940 Hi-C contacts, and 2,940,433,604 Hi-C read pairs in stimulated T cells yielding 1,853,522,064 Hi-C contacts. Loci were assigned to A and B compartments at 500 kB resolution. Standard loops were annotated using HiCCUPS at 5kB, 10kB, and 25kB resolutions with default Juicer parameters. This yielded a list of 2,640 loops in unstimulated T cells and 4,614 loops in stimulated T cells at MAPQ > 30. Differential loop calling with HiCCUPSDiff at 5kB, 10kB, and 25kB identified 21 loops as being significantly enriched for unsimulated T cells and 334 loops as being significantly enriched for simulated T cells. Contact domains were annotated using the Arrowhead algorithm with default Juicer parameters at 10kB resolution for unstimulated T cells and 5kB for stimulated T cells. This yielded a list of 1,549 domains in unstimulated T cells and 4,008 domains in stimulated T cells at MAPQ > 30. We also ran Arrowhead with these same respective parameters on MAPQ > 0 Hi-C maps, which yielded a list of 1,810 domains in unstimulated T cells and 4,419 domains in stimulated T cells. The Hi-C maps and feature annotations were visualized using the Juicebox software^71^.

### Determination of distance from ATAC-peak to contact domains

We determined the distance from each ATAC-peak to the middle of the closest contact domain. We analyzed the following features: (**1**) all ATAC-peaks; (**2**) *local -*ATAC-peaks; and all ATAC-peaks and *local -*ATAC-peaks containing (**3**) BATF, (**4**) ETS1, or (**5**) CTCF motifs. We normalized the distances from each peak to the closest domain by the length of the domain. In order to determine that the distribution of the distance between a given peak and a contact domain is different than the null distribution, we kept the length of each contact domain constant and shuffled the positions of the contact domain. The distances from each peak to the contact domain were binned into 30 bins and divided by the binned distances between a given peak and the shuffled contact domains to determine enrichment at each position.

### Co-accessible peak analysis

To identify co-accessible peaks, we tested for correlation between every pair of the 167,140 ATAC-peaks within 1.5 Mb of each other using a linear regression in Matrix eQTL^72^. We first normalized the ATAC-peaks by (**1**) removing sequencing depth bias using a median normalization, (**2**) standardizing the matrix by subtracting out the mean and dividing by the standard deviation for each peak; and (**3**) quantile normalization of the matrix^73^. Adjusting for sex and 15 principle components, we used Matrix eQTL to identify 2,158 pairs of co-accessible peaks (1,809 unique ATAC-peaks, FDR < 0.05).

### RNA-seq analysis

25bp paired end RNA-seq reads were aligned to the hg19 using UCSC transcriptome annotations. Expression levels (expected counts) were determined using RSEM^74^. We applied TMM normalization to the expected counts using the edgeR package and kept genes that had TMM count > 1 in at least 75% of the samples. For the mapping of eQTNs, we inputted expected counts for filtered genes into RASQUAL^43^, which performs internal normalization. For the heritability analyses, we used log-transformed TMM counts of filtered genes in order to fit linear mixed models.

### Percentage of GWAS loci overlapping

The GREGOR suite^57^ was used for calculating the percentage of GWAS loci in features of interest: (**1**) peaks differentially accessible in Th and Th_stim_ cells; (**2**) co-accessible peaks, and (**3**) *local*-ATAC-peaks. For *local*-ATAC-peaks, peaks were randomly permuted, while retaining the width of each peak to assess the expected GWAS enrichment.

### Percentage overlapping T cell annotations

Using the Homer suite annotatePeaks.pl with the –genomeOntology option, we calculated how many of the Th_stim_-specific peaks, co-accessible peaks and *local*-ATAC-peaks fall T cell enhancers^18^. For co-accessible peaks and *local*-ATAC-peaks peak subsets, peaks were randomly permuted, while retaining the width of each peak to assess the expected T cell enrichment.

### Proportion in super-enhancer regions

Using the BEDtools intersect function, we calculated how many of the co-accessible peaks, co-accessible *local*-ATAC-peaks, and co-accessible *local*-ATAC-peaks that fall in Hi-C domains are also in stimulated Th super-enhancers (as reported in Hinsz et al.^41^). Super enhancers were shuffled by randomly sampling the genome to the same number of features and retaining the length of each super enhancer to generate a background.

### Proportion co-accessible peaks in known regulatory elements

Using the BEDtools intersect function, we annotated each peak in our unique pairs of co-accessible peaks as residing in a known Th stim super enhancer (as reported in Hinsz et al.^41^), promoter (as reported in Fahr et al.^18^), and T cell promoter (as reported in Fahr et al.^18^). We determined if each peak in a pair of peaks resided in a promoter and a promoter, promoter and a super enhancer, a promoter and an enhancer, an enhancer and an enhancer, an enhancer and a super enhancer, and a super enhancer and a super enhancer. Finally, we calculated the proportion of pairs of co-accessible peaks that were annotated as each pair regulatory elements. As background (‘non co-accessible peak’) we used pairs of ATAC-peaks with P-value > 0.9, sampled to the same number as co-accessible peaks, and performed the same analysis.

### Gkm-SVM and deltaSVM

We ran gkm-SVM^75,76^ on 24,745 300bp ATAC-peaks centered on MACS summits using default parameters and an equal size GC matched negative set, excluding from training any region containing a SNP to be scored by deltaSVM, and repeated with 5 independent negative sets, and averaged the deltaSVM predictions, as previously described^45^. We then calculated deltaSVM for each SNP in a *local*-ATAC-peak p-value < 10^-5^, scoring 903 SNPs in 888 loci. We find a Pearson correlation of R=0.627 between ATAC-QTN beta and the largest deltaSVM SNP. 777 of the peak p-value SNPs had the largest deltaSVM, but 111 flanking SNPs scored more highly than the peak p-value SNP and disrupt immune associated TF binding sites. While the gkm-SVM weights fully specify the deltaSVM score, for interpretation we associated the large gkm-SVM weights with the most similar TF PWM from a catalog of JASPAR, Transfac, Uniprobe, and Homer motifs.

### Heritability of gene expression and ATAC-peaks

Data for the heritability analysis of gene expression and ATAC-peaks was prepared in the following way. Each ATAC-peak was residualized against its 10 principal components and sex while the expression of each gene was residualized for 12 principal components and sex.

For the univariate analyses, restricted maximum likelihood heritability (h^2^) estimates were calculated using GCTA software^55^ with algorithm 1 and no constraints on heritability (*i.e*., h^2^ can be less than 0), while the bivariate analysis was run constrained. For the gene expression heritability and ATAC-peak heritability analyses, we used genotypes +/- 500 kb from the transcription start site of the gene and center of each ATAC-peak, respectively. Of the 64,188 ATAC-peaks and 3,318 *local*-ATAC-peaks, 32,317 and 2,444 converged respectively. The bivariate GCTA analysis used genotypes +/-500kb from the transcription start site of the gene. Randomly sampled ATAC-peaks (non *local*-ATAC-peaks) and permuted ATAC-peaks were plotted as background at the same number of the tested *local*-ATAC-peaks (N=161, standard errors < 1).

## Acknowledgements

We thank the ImmVar participants. We would like to thank Jason Buenrostro for critical reading of the manuscript and advice on ATAC-seq analysis, Jenna Pfiffner and Charles Fulco for initial experimental help with ATAC-seq, Alicia Schep for ATAC-seq nucleosome free caller, Natasha Asinovski and Ho-keun Kwon for help setting up primary T cell cultures and members of the Regev laboratory for discussions. M.B. and K.L.H. are supported by NIH HG007348 to M.B., H.Y.C. is supported by NIH grant P50-HG007735, C.S.C is supported by the NIH through a Ruth L. Kirschstein National Research Service Award (F32-DK096822). This work was supported by the Klarman Cell Observatory at the Broad Institute. A.R. is a Howard Hughes Medical Institute Investigator. Raw data are deposited to the Gene Expression Omnibus with accession no. GSE86888.

